# Manipulation of neuronal activity in the entorhinal-hippocampal circuit affects intraneuronal amyloid-β levels

**DOI:** 10.1101/2022.07.05.498797

**Authors:** Christiana Bjorkli, Nora C Ebbesen, Joshua B. Julian, Menno P Witter, Axel Sandvig, Ioanna Sandvig

## Abstract

One of the neuropathological hallmarks of Alzheimer’s disease (AD) is the accumulation of amyloid-β (Aβ) plaques, which is preceded by intraneuronal build-up of toxic, aggregated Aβ during disease progression. Aβ plaques are first deposited in the neocortex before appearing in the medial temporal lobe, and tau pathology with subsequent neurodegeneration in the latter anatomical region causes early memory impairments in patients. Current research suggests that early intraneuronal Aβ build-up may begin in superficial layers of lateral entorhinal cortex (LEC). To examine whether manipulation of neuronal activity of LEC layer II neurons affected intraneuronal Aβ levels in LEC and in downstream perforant path terminals in the hippocampus (HPC), we used a chemogenetic approach to selectively and chronically silence superficial LEC neurons in young and aged 3xTg AD mice and monitored its effect on intraneuronal Aβ levels in LEC and HPC. Chronic chemogenetic silencing of LEC neurons led to reduced early intraneuronal Aβ in LEC and in projection terminals in the HPC, compared with controls. Early intraneuronal Aβ levels in the downstream HPC correlated with activity levels in superficial layers of LEC, with the subiculum being the earliest subregion involved, and our findings give evidence to early AD neuropathology originating in select neuronal populations.

## 1. Introduction

Alzheimer’s disease (AD) constitutes most dementia cases, and knowledge is still lacking regarding the underlying factors that lead to the disease. The accumulation of hyperphosphorylated, misfolded tau proteins into neurofibrillary tangles (NFTs), coupled with deposition of amyloid-beta (Aβ) into extracellular plaques, are the two hallmark pathological features of AD in the brain [1-4]. The pathophysiology of AD is marked by slowly progressing abiotrophic neurodegeneration, which begins in the medial temporal lobes (MTLs), a region known to be critical for learning and memory [5, 6]. The MTLs comprise the hippocampus (HPC) along with the surrounding hippocampal region, including the entorhinal cortex (EC) that serves as the primary interface for HPC-neocortex circuits [5, 7, 8].

AD neuropathology follows a hierarchical deposition pattern in anatomically and functionally connected brain regions [3, 4]. In patients, the lateral EC (LEC) is an early region of neuronal loss [9] and likely the brain area from where AD pathology invades other brain regions [10, 11]. Neurons in the superficial layers of LEC form synapses via the perforant pathway with all hippocampal subregions, including the dentate gyrus (DG), cornu ammonis (CA) 1 and 3, and subiculum (Sub). The encoding of various forms of memory [12, 13], but particularly contextual memory [14], requires an intact EC-HPC circuit. It has remained elusive where toxic intraneuronal Aβ originates in the brain, but current research suggests that this build-up may occur in superficial layers of EC during early stages of AD [11, 15, 16].

The presence of soluble, oligomeric intraneuronal Aβ is considered more neurotoxic compared to the later-developed Aβ plaques during progression of AD [17-26]. Since the EC-HPC circuit is affected [27-29], and intraneuronal Aβ is present in EC during early stages of the disease, changes originating in EC could affect neuropathological development in downstream perforant path terminals [30-34]. In this study we chronically silenced neuronal activity of LEC layer II neurons of young and aged 3xTg AD mice, by locally expressing inhibitory designer receptors exclusively activated by designer drugs (DREADDs) in these neurons. We subsequently activated the DREADDs by local infusions of the novel ligand deschloroclozapine (DCZ) and examined its effect on early intraneuronal Aβ build-up in LEC layer II and downstream HPC by immunolabelling.

## 2. Methods

### 2.1. Animals

3xTg AD mice (MMRRC Strain #034830-JAX; RRID: MMRRC_034830-MU; *n* = 27) and control B6129 mice (Strain #:101045; RRID:IMSR_JAX:101045; *n* = 17) were included in these experiments. 3xTg AD mice contain human transgenes for amyloid precursor protein (*APP*) bearing the Swedish mutation, presenilin-1 (*PSEN1*) containing an M146V mutation, and microtubule-associated protein tau (*MAPT*) containing an P301L mutation. The donating investigators of the 3xTg AD mouse model has previously communicated to Jackson Laboratories that male transgenic mice may not exhibit all phenotypic traits of AD [35]. Therefore, only female mice were included in these experiments. All housing and breeding of animals was approved by the Norwegian Animal Research Authority and is in accordance with the Norwegian Animal Welfare Act §§ 1-28, the Norwegian Regulations of Animal Research §§ 1-26, and the European Convention for the Protection of Vertebrate Animals used for Experimental and Other Scientific Purposes (FOTS ID 21061). The animals were kept on a 12h light/dark cycle under standard laboratory conditions (19-22°C, 50-60% humidity), and had free access to food and water.

### 2.2. Stereotactic viral injections

Mice were anaesthetized with isoflurane (4% induction and 1-3% maintenance, IsoFLo vet., Abbott Laboratories, Chicago IL, USA) and oxygen (0.3 - 0.4 %) in an induction chamber before being transferred to a stereotaxic frame (David Kopf Instruments, California, USA) and placed on a temperature-controlled heating pad (40.5 – 41 °C). Mice were administered analgesics (Temgesic, Invidor UK, Slough, Great Britain [0.05-0.09 mg/kg], and Metacam, Boehringer Ingelheim Vetmedica, Copenhagen, Denmark [0.05-0.15 mg/kg]) and a local anesthetic (Marcain, Aspen Pharma, Ballerup, Denmark [0.03-0.18 mg/kg]) subcutaneously, and an incision was made to expose the skull. To horizontally align the skull, the ventral distance down to the stereotaxic landmarks bregma and lambda were assessed, and the medial-lateral alignment of the skull was examined by comparing the ventral distance down to the stereotaxic landmarks ±1 mm laterally from the midline of the brain. A craniotomy was made at 0.5 mm anterior to lambda and ∼4 mm lateral (dependent on animal weight) to the midline. A Hamilton microsyringe (Neuros 32-gauge syringe, 5 μl, Hamilton company, Nevada, USA) was lowered vertically into the brain to a depth ∼3.6 mm (dependent on animal weight) from the surface, and 500-1000 nl of viruses was injected using a microinjector (Nanoliter 2010, World Precision Instruments Inc., USA). We used the following adeno-associated viruses (AAVs) for injections: AAV8 human M4 muscarinic (hM4) inhibitory DREADDs (hM4D_i_)-mCherry (referred to as ‘AAV-hM4D_i_’ hereafter) and AAV8 mCherry (referred to as ‘AAV-mCherry’ hereafter), both with the calcium/calmodulin-dependent protein kinase (CaMKII) promoter to drive expression of the DREADD in excitatory cells only). AAVs were generated by Dr. Nair at the Viral Vector Core Facility, at the Kavli Institute for Systems Neuroscience, Trondheim, Norway. AAV-mCherry was injected into the contralateral hemisphere of the one injected with AAV-hM4D_i_ in a portion of the mice (*n* = 14). The microsyringe was kept in place for 10 minutes prior to and after the injection, to minimize potential upward leakage of the viral solution. Metacam was given within 24 hours post-surgery. Animals were implanted with osmotic minipumps 6-7 days following injections.

### 2.3. Activation of inhibitory designer receptors

An Alzet osmotic minipump (model 1004, flow rate: 0.11 μl/hour, Durect Corporation, California, USA) was primed with 100 μl DCZ (MedChemExpress, USA) [36] at a dosage of 100 μg/kg (or with sterile saline for controls), at 38°C in sterile saline for 48 hours prior to implantation.

Following the same procedure as the viral injection surgery up until the target coordinate was derived, mice were implanted with minipumps subcutaneously on the flank, slightly posterior to the scapulae. A hemostat was inserted into the incision to make a pocket for the minipump large enough to allow free movement of the animal (approximately 1 cm longer than the minipump). For the implantation of the intracranial cannula attached to the minipump (Durect Corporation, California, USA) a craniotomy was made at 0.1 mm posterior to bregma and 1.2 mm lateral to the midline to target the lateral ventricle. The intraventricular cannula was lowered into the brain to a depth 2.75 mm from the surface of the brain, attached to the skull using superglue and dental cement (Dentalon Plus, Cliniclands AB, Trelleborg, Sweden), while the catheter and minipump was secured under the skin with sutures. Analgesics were repeated within 24 hours post-surgery.

### 2.4. Context-dependent spatial memory testing

To confirm functional silencing of LEC layer II, we examined context-dependent spatial memory (**Supplementary Fig. 1**) in mice with intraventricular infusions of DCZ or saline. The basic training and testing protocol is also described in [37]. But in brief, disoriented mice were initially trained to dig for buried food rewards (Weetos choco, Nestlé S.A., Vevey, Switzerland) in two different chambers, one with square boundaries (4 × 29.25 cm) and one with circle boundaries (157 cm circumference). All chambers were built out of rectangular Legos (2 × 1 cm; Lego A/S, Billund, Denmark), and were 15 cm tall. Rewards were buried under odor-scented bedding in cups embedded in the chamber floors; the odor-masked bedding consisted of 1 g of ground fresh ginger for every 100 g of bedding. Each chamber was surrounded by the same distal cues for orientation. There were four possible reward locations in each chamber, and the rewarded location differed between the square- and circle-chamber relative to the common reference frame provided by the distal cues. The training phase consisted of four training trials per chamber per day for 2 days, with successive trials alternated across chambers (8 trials total in the square chamber and 8 trials total in the circle chamber). If a given mouse achieved 66.6% correct performance during training, contextual memory was then tested in 4 testing sessions across 4 days, with 8 trials per session. Four 3xTg AD mice with silenced LEC were unable to complete the training phase and were therefore excluded from further analyses. During each testing session, the first two trials consisted of spatial memory being tested in the square- and circle-chamber with rewards. In trials 3-6, spatial memory was tested in four chambers with morphed boundary geometry, which continuously ranged from most-square-like to most-circle-like: a pentagon (5 × 31.4 cm), a hexagon (6 × 26.16 cm), an octagon (8 × 19.6 cm), and a decagon (10 × 15.7 cm). During the final 2 trials of each testing session (trials 7-8), the animals were again tested in the square and circle chambers with rewards. The order of the square-, circle- and morphed-chambers across trials in each session was randomized but was the same for each animal on a given day’s session.

Dig locations and time spent in these locations were calculated using ANY-maze video tracking system (Stoelting Europe) via an overhead, centrally located camera (DMK 22AUC03 USB 2.0 monochrome industrial camera, The imaging Source Europe, Germany). Animals had access to water ad libitum but were maintained at 90-95 % of their free-feed weight to increase task motivation.

### 2.5. Tissue processing

Animals were sacrificed after neuronal silencing and/or behavioral testing. The animals were subjected to a novel object (Duplo Lego) 90 min prior to perfusions [38]. Mice were administered a lethal dose of sodium pentobarbital and transcardially perfused with Ringer’s solution followed by paraformaldehyde (PFA, 4 %, Merck Millipore, Darmstadt, Germany) in 125 mM phosphate buffer (PB). Brains were extracted and fixed for a minimum of 24 hours in PFA at 4 °C and transferred to a 2 % dimethyl sulfoxide solution (VWR International, Radnor, PA, USA) prepared in PB for at least 24 hours at 4 °C. Brains were sectioned coronally at 40 µm on a freezing sliding-microtome, and an incision was made in the contralateral hemisphere of the one injected with the experimental virus to mark the control hemisphere.

### 2.6. Immunohistochemical processing

Immunolabeling for Aβ, reelin, mCherry, hM4 and cFos was conducted on tissue from all animals. See [39-42] for detailed protocols, and **Supplementary Table 1** for details regarding the antibodies used. Sections were scanned using a Mirax-midi slide scanner (objective 20X, NA 0.8; Carl Zeiss Microscopy, Oberkochen, Germany), using either reflected fluorescence (for sections stained with a fluorophore) or transmitted white light (for sections stained with Cresyl violet) as the light source.

### 2.7. Quantification of cFos+ and amyloid-β+ neurons

Series of sections were chosen randomly and coded to ensure blinding to the investigators. The number of cells containing cFos and intraneuronal Aβ in LEC and HPC were estimated with Ilastik [43] using the Density Cell Counting workflow. LEC and HPC was delineated using cytoarchitectonic features in sections stained with Nissl, based on The Paxinos & Franklin Mouse Brain Atlas [44]. The regions constituting the human MTLs are equivalent to the hippocampal region in rodents, consisting of the HPC (CA1-3, DG, and Sub) and the adjacent perirhinal, entorhinal, and parahippocampal cortices [45]. The same surface area and rostrocaudal levels of each brain region was selected, and at least 4 brain sections containing infused hemispheres were used for quantification. Damaged regions of brain sections were excluded from analyses to avoid false-positive antibody expression.

### 2.8. Statistics

Effect size (Cohen’s D) was calculated based on initial experiments between animals injected with experimental and control viruses and the resulting difference in LEC layer II cFos+ cells. Based on each group consisting of at least 4 animals, a large effect size of 1.57 was calculated [46]. Most of the dataset displayed a normal distribution (Shapiro-Wilk test) and therefore two-tailed, unpaired t-tests were used to compare mean differences. For the minor parts of the dataset that was not normally distributed, nonparametric statistical tests were used to compare means (Mann-Whitney U). All reported statistics were based on two-tailed significance tests. Statistical comparisons of behavioral data across LEC-silenced and control mice were conducted based on trial-wise pooling of data across mice separately for each group. Behavioral performance in the morphed environments of the contextual memory task was calculated as follows: dig in square-consistent location = 1, dig in circle-consistent location = 0, dig in any other location = 0.5. Context-consistency of reward locations was determined relative to the common reference frame defined by the distal cues shared across all contexts. We then assessed whether performance in the morphed environments was associated with more context-appropriate choices for control animals treated with a vehicle compared to LEC-silenced animals. All statistical tests and graphs were made in Prism 9 (GraphPad Software Inc., CA, USA).

## 3. Results

### 3.1. Verification of chronic neuronal silencing of lateral entorhinal cortex layer II at the molecular and functional level

We first verified neuronal silencing following injections of AAV-hM4D_i_ in LEC layer II and intraventricular DCZ infusions for 2-3 weeks (**Fig. 1A, B; Supplementary Fig. 2A**). Our injections were limited to reelin-expressing principal neurons in LEC layer II (**Supplementary Fig. 2B**). Moreover, hM4 receptor labelling was present in LEC layer II neurons infected with AAV-hM4D_i_, but not when infected with the control virus AAV-mCherry (**Supplementary Fig. 2C**). hM4 receptor labelling was also not present in LEC layer II neurons infected with AAV-hM4D_i_ after saline infusions (**Supplementary Fig. 2D**). After injections of AAV-hM4D_i_, we observed reduced cFos expression in LEC layer II of 3xTg AD mice after DCZ (but not saline) infusions (t = 19.43, p < .05, unpaired two-tailed t-test; **Fig. 1C-E**).

**Figure 1.**
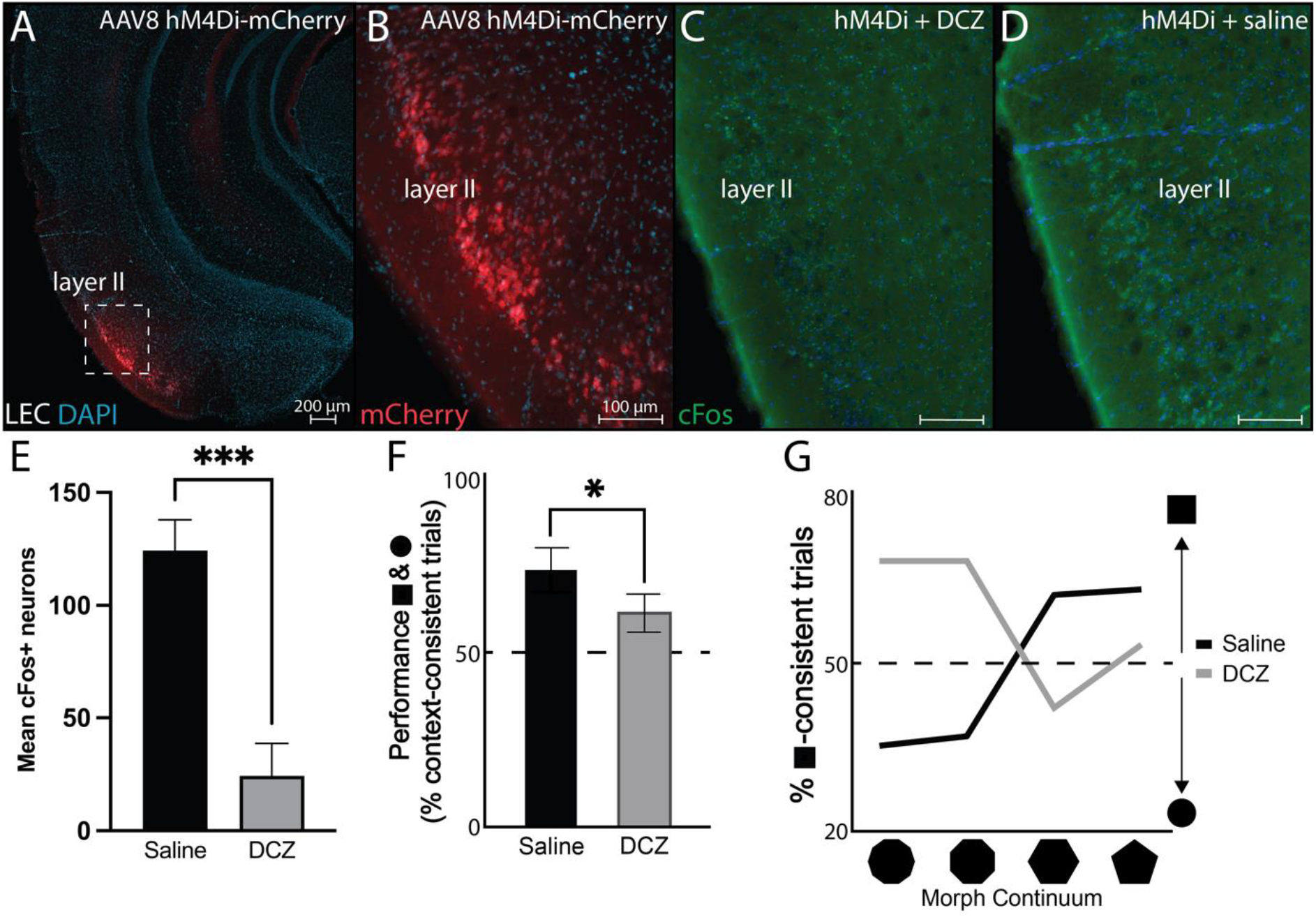
Chronic silencing of LEC layer II attenuates neuronal activity. **A)** AAV-hM4D_i_ (mCherry; red) injection location in LEC layer II of 3xTg AD mice. **B)** Higher magnification mCherry+ (red) LEC layer II neurons in 3xTg AD mice. **C)** cFos+ (green) LEC layer neurons following AAV-hM4D_i_ injections in LEC layer II and intraventricular infusions of DCZ for 2 weeks in 3xTg AD mice. **D)** cFos+ (green) LEC layer neurons following injections in LEC layer II and intraventricular infusions of saline for 2 weeks in 3xTg AD mice. **E)** Mean cFos+ LEC layer II neurons in infused and non-infused hemispheres (<4 brain sections) after AAV-hM4D_i_ injections and DCZ (*n* = 25) or saline (*n* = 9) intraventricular infusions. **F)** Mean behavioral performance (% of trials with context-appropriate choices) in 3xTg AD mice with hM4D_i_ in LEC layer II and DCZ (*n* = 2) or saline (*n* = 2) infusions in the square and the circle contexts. **G)** Percentage of trials with square context-consistent digs in 3xTg AD mice with hM4D_i_ in LEC layer II and DCZ (*n* = 2) or saline (*n* = 2) infusions in each of the four morphed environments, ranging from most circle-like to most square-like in boundary geometry shape. Abbreviations; AAV: adeno-associated virus; hM4D_i_: human M4 muscarinic (hM4) inhibitory DREADDs; LEC: lateral entorhinal cortex; DCZ: deschloroclozapine. Error bars denote ±1 SD in panel E, and ±1 SEM in panels F-G; * p < .05, *** p < .001.

To confirm functional silencing of LEC layer II, we examined context-dependent spatial memory (**Supplementary Fig. 1**). Mice were trained to dig for buried food rewards in two different contexts, one square and one circle. Mice with silenced LEC initially searched in context-appropriate reward locations less often than controls (p < .05; Fischer’s exact test; **Fig. 1F**). To confirm that this spatial memory impairment was due to disrupted contextual processing, we further examined contextual memory recall in environments with progressive morphs of boundary geometry, increasing the number of boundary walls from most square-like to most circle-like. This type of pattern completion dependent behavior is thought to depend on the normal function of the HPC [47, 48]. Compared to controls, mice infused with DCZ searched less often in context-consistent reward locations (χ2 = 24.13, p < 10^−5^; Pearson’s Chi-squared test; **Fig. 1G**). These effects are unlikely to be explained by domain general search behavior, as response latency and movement speed did not differ between DCZ mice and controls (data not shown). Thus, silencing of LEC layer II neurons led to behavioral changes associated with a dysfunctional EC-HPC circuitry [27-29].

### 3.2. Chronic neuronal silencing of lateral entorhinal cortex layer II lowered intraneuronal amyloid-β levels

Principal cells in LEC layer II can be distinguished from nearby neuronal populations based on their expression of the extracellular matrix glycoprotein reelin (**Fig. 2A**) [7]. Here we observed that reelin was expressed more in dorsal compared to ventral portions of LEC layer II (t = 9.43, p < .001, unpaired two-tailed t-test; **Fig. 2A, D**). Previous research suggests a link between reelin+ LEC layer II neurons and the build-up of toxic intraneuronal Aβ [10, 34], and we observed that intraneuronal Aβ colocalized with reelin, and was similarly more abundant in dorsal compared to ventral LEC layer II neurons (t = 6.86, p < .0001, unpaired two-tailed t-test; **Fig. 2B, C, E**). Therefore, we next examined whether LEC layer II neuronal silencing affected intraneuronal Aβ build-up in the same population of neurons. We observed less intraneuronal Aβ in LEC layer II neurons infected with AAV-hM4D_i_ and infused with DCZ than when infused with saline (t = 3.52, p < .01, unpaired two-tailed t-test; **Fig. 2F-H**), but no change in reelin levels of these same neurons (**Supplementary Fig. 3A**). Within 3xTg AD mice with silenced LEC layer II, we observed reduced intraneuronal Aβ in LEC layer II neurons to a greater degree in the hemisphere injected with AAV-hM4D_i_ compared to the contralateral non-injected/injected with AAV-mCherry hemisphere (**Supplementary Fig. 3B**).

**Figure 2.**
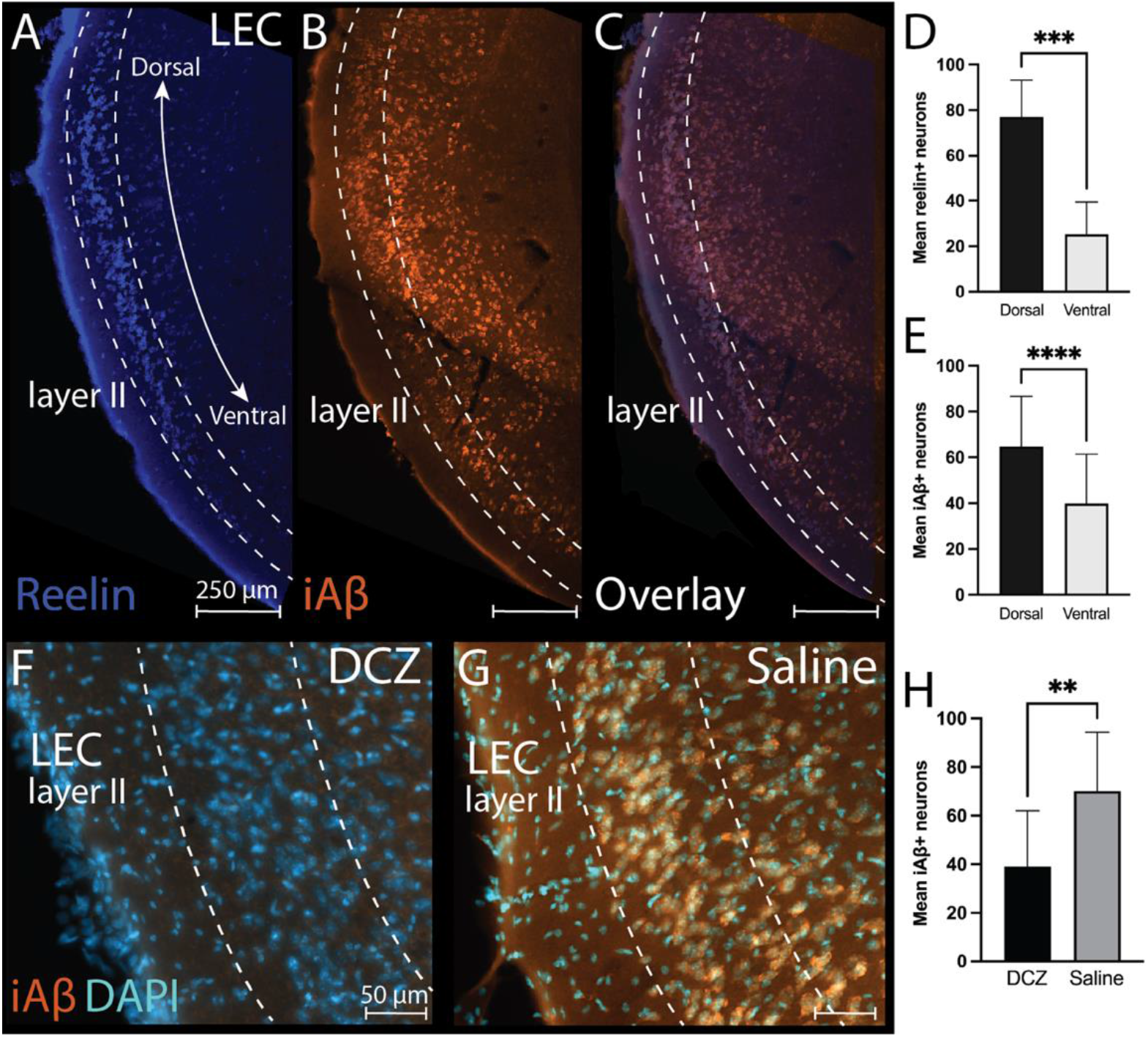
Targeted silencing of reelin-expressing LEC layer II results in reduced intraneuronal Aβ levels. **A)** Expression of reelin (blue) in the dorsal-ventral gradient of LEC layer II neurons. **B)** Expression of intraneuronal Aβ (McSA1 antibody; orange) in the dorsal-ventral gradient of LEC layer II neurons. **C)** Co-expression of reelin (blue) and intraneuronal Aβ (orange) in the dorsal-ventral gradient of LEC layer II neurons. **D)** Bar graph displaying mean reelin+ LEC layer II neurons in dorsal and ventral LEC layer II (<4 brain sections). **E)** Bar graph displaying mean intraneuronal Aβ+ LEC layer II neurons in dorsal and ventral LEC layer II (<4 brain sections). **F)** Expression of intraneuronal Aβ (orange) in LEC layer II hM4D_i_-infected neurons following DCZ infusions (*n* = 25). **G)** Expression of intraneuronal Aβ (orange) in LEC layer II hM4D_i_-infected neurons following saline infusions (*n* = 9). **H)** Bar graph displaying mean intraneuronal Aβ+ LEC layer II neurons in infused and non-infused hemispheres (<4 brain sections) after injections of AAV-hM4D_i_ and DCZ (*n* = 25) or saline (*n* = 9) intraventricular infusions. Abbreviations; LEC: lateral entorhinal cortex; iAβ: intraneuronal amyloid-β; DCZ: deschloroclozapine. Error bars denote ±1 SD; ** p < .01, *** p < .001, **** p <.0001.

### 3.3. Chronic silencing of lateral entorhinal cortex layer II lowered intraneuronal amyloid-β levels in hippocampal terminal fields

We next tested whether chronic LEC layer II neuronal silencing reduced early intraneuronal Aβ presence in its projection terminals in HPC, specifically in those subregions of the HPC known to exhibit Aβ pathology in patients (i.e., dSub, CA1 and Sub) [4]. Chronic LEC layer II neuronal silencing reduced intraneuronal Aβ in projection terminals in the HPC broadly (t = 2.87, p < .01, unpaired two-tailed t-test; **Fig. 3A-E**). As expected, injections of AAV-hM4D_i_ into LEC layer II followed by saline infusions did not cause a reduction of intraneuronal Aβ in the HPC (n.s.; **Fig. 3F-J**). We also found no change in intraneuronal Aβ levels in downstream HPC when the length of inhibition was 2 or 3 weeks (n.s.; **Supplementary Fig. 3C**). Within HPC of young 3xTg AD mice (2-months-old), intraneuronal Aβ was reduced in all subregions after LEC layer II silencing, with the greatest reduction observed in ventral Sub (t = 3.99, p < .01, unpaired two-tailed t-test; **Fig. 3D, K**). In older 3xTg AD mice (4- and 6-months-old), intraneuronal Aβ was reduced in all subregions after silencing, but CA1 (4-months-old: t = 2.72, p < .05, unpaired two-tailed t-test; **Fig. 3A-C**) and dorsal Sub (4-months-old: t = 2.67, p < .05; 6-months-old: t = 3.08, p < .05, unpaired two-tailed t-tests; **Fig. 3A-C**) displayed greater reduction (**Fig. 3L, M**).

**Figure 3.**
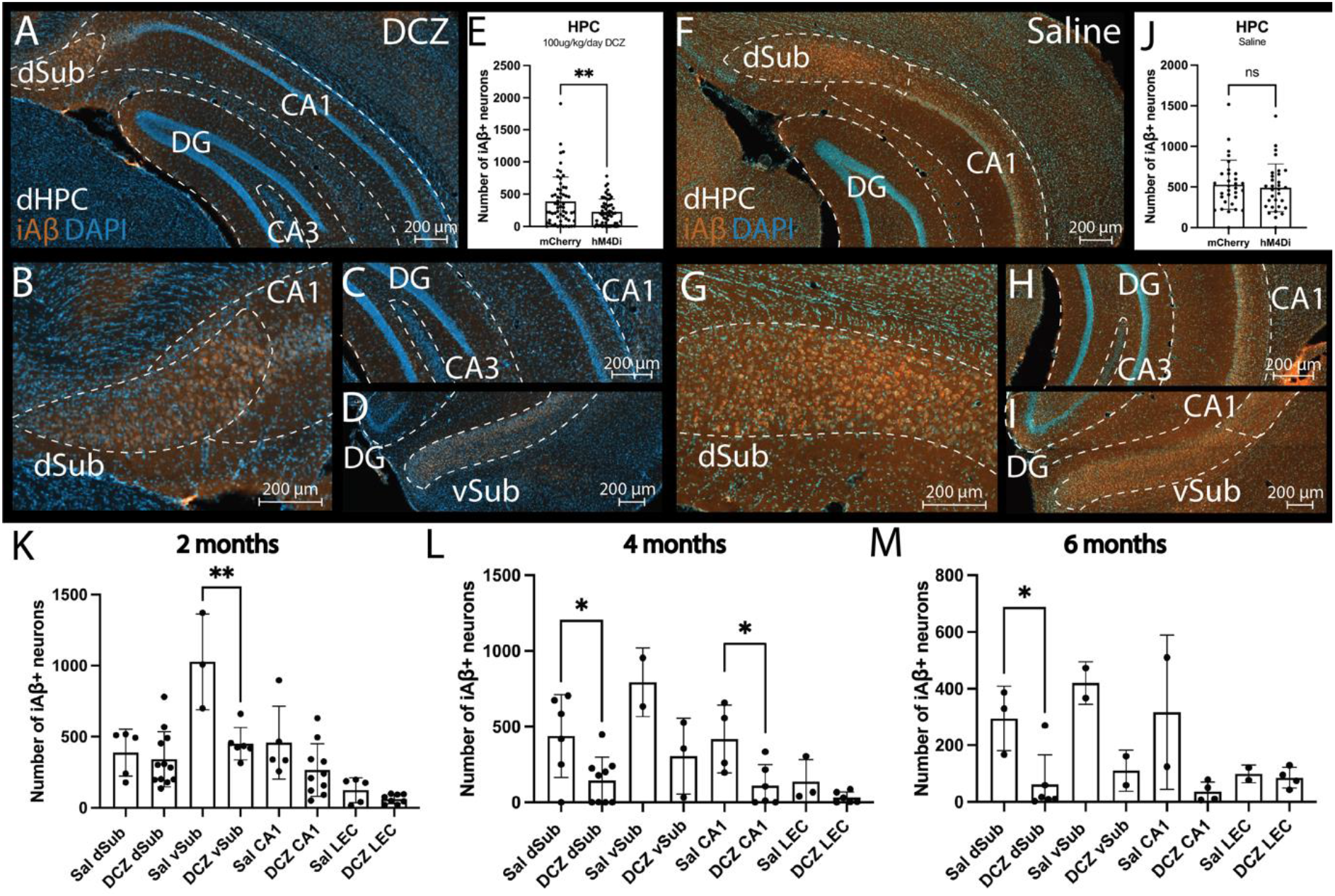
Reduced expression of intraneuronal Aβ in downstream HPC following LEC layer II neuronal silencing. **A)** Macro-view of the expression of intraneuronal Aβ (McSA1 antibody; orange) in the dorsal HPC of 3xTg AD mice with silenced LEC layer II and DCZ infusions. **B)** Higher magnification view of intraneuronal Aβ expression (orange) in dorsal Sub and CA1 of 3xTg AD mice with silenced LEC layer II and DCZ infusions. **C)** Higher magnification view of intraneuronal Aβ expression (orange) in CA1/3 and DG of 3xTg AD mice with silenced LEC layer II and DCZ infusions. **D)** Higher magnification view of intraneuronal Aβ expression (orange) in ventral Sub of 3xTg AD mice with silenced LEC layer II and DCZ infusions. **E)** Box plots showing mean intraneuronal Aβ+ neurons in HPC following injections of either AAV-hM4D_i_ (*n* = 25) or AAV8 mCherry (*n* = 8) with DCZ infusions. Each dot represents a brain section containing the hippocampal subregions, and sections spanned the same rostro-caudal level for both AAV-hM4D_i_ and AAV-mCherry injections. **F)** Macro-view of the expression of intraneuronal Aβ (McSA1 antibody; orange) in the dorsal HPC of 3xTg AD mice with LEC layer II AAV-hM4D_i_ injections and saline infusions. **G)** Higher magnification view of intraneuronal Aβ expression (orange) in dorsal Sub of 3xTg AD mice with LEC layer II AAV-hM4D_i_ injections and saline infusions. **H)** Higher magnification view of intraneuronal Aβ expression (orange) in CA1/3 and DG of 3xTg AD mice with LEC layer II AAV-hM4D_i_ injections and saline infusions. **I)** Higher magnification view of intraneuronal Aβ expression (orange) in CA1 and ventral Sub of 3xTg AD mice with LEC layer II AAV-hM4D_i_ injections and saline infusions. **J)** Box plots showing mean intraneuronal Aβ+ neurons in HPC following injections of either AAV-hM4D_i_ (*n* = 9) or AAV8 mCherry (*n* = 6) with saline infusions. Each dot represents a brain section containing the hippocampal subregions, and sections spanned the same rostro-caudal level for both AAV-hM4D_i_ and AAV-mCherry injections. **K)** Box plots showing mean intraneuronal Aβ+ neurons in hippocampal subregions (dSub, vSub, CA1) and LEC in 2-month-old 3xTg AD mice after AAV-hM4D_i_ injections and DCZ (*n* = 8) or saline (*n* = 2) infusions. **L)** Box plots showing mean intraneuronal Aβ+ neurons in hippocampal subregions (dSub, vSub, CA1) and LEC in 4-month-old 3xTg AD mice after AAV-hM4D_i_ injections and DCZ (*n* = 11) or saline (*n* = 3) infusions. **M)** Box plots showing mean intraneuronal Aβ+ neurons in hippocampal subregions (dSub, vSub, CA1) and LEC in 6-month-old 3xTg AD mice after AAV-hM4D_i_ injections and DCZ (*n* = 5) or saline (*n* = 2) infusions. Abbreviations; dSub: dorsal subiculum; DG: dentate gyrus; CA: cornu ammonis; iAβ: intraneuronal amyloid-β; dHPC: dorsal hippocampus; DCZ: deschloroclozapine; hM4D_i_: human M4 muscarinic (hM4) inhibitory DREADDs; vSub: ventral subiculum; LEC: lateral entorhinal cortex; sal: saline. Error bars denote ±1 SD; ** p <.01; ; * p <.05; ns: non-significant.

## 4. Discussion

To examine whether intraneuronal Aβ build-up correlated with neuronal activity levels, we silenced LEC layer II neuronal activity in 3xTg AD mice. This was done by targeted AAV-hM4D_i_ injections into LEC layer II followed by intraventricular infusions of the novel DREADD ligand DCZ. We opted to use DCZ rather than the conventionally used clozapine N-oxide (CNO) ligand, since the former ligand does not convert to clozapine and has a higher affinity to DREADDs [36]. Neuronal silencing caused context-dependent spatial memory deficits, indicating a functional role of LEC layer II neurons in cognitive functions known to be impaired early during AD progression in patients. Moreover, chronic inhibition of LEC layer II caused a reduction of intraneuronal Aβ levels within LEC neurons. Critically, chronic LEC layer II neuronal silencing also caused a reduction of intraneuronal Aβ levels in the subregions CA1 and Sub of the HPC, suggesting that activity levels in LEC affected intraneuronal Aβ in projections terminals in the HPC. The first subregion of the HPC that displayed reduced intraneuronal Aβ levels after neuronal silencing was the Sub. Together, these results suggest that intraneuronal Aβ build-up in LEC layer II and its perforant path terminals is regulated by neuronal activity.

It has previously been suggested that toxic intraneuronal Aβ build-up can appear in multiple brain regions simultaneously, before it encompasses the majority of association cortex [3, 4, 49, 50]. Alternatively, toxic Aβ may propagate in a cell-to-cell manner, consistent with the observation that the protein is found intracellularly [17, 51, 52]. However, neuroimaging and autopsy studies of AD patients have not pinpointed a precise anatomical origin of intraneuronal Aβ build-up. Animal studies have begun to shed light on the potential origin of early intraneuronal Aβ build-up in the brain. Specifically, it has been shown that *APP* mutations, of which can cause increased Aβ production [53], can be transported anterogradely via perforant path projections from superficial layers of EC to DG in rats [54], resulting in high levels of soluble Aβ peptides and plaque deposits in DG [55]. Recent findings suggest that lowered levels of reelin in LEC layer II principal neurons led to reduced intraneuronal Aβ in these neurons, as well as in projection terminals in the HPC [10]. This is in line with our current findings, which provide evidence of activity-dependent changes in intraneuronal Aβ build-up in the EC-HPC circuit.

In AD patients, extracellular Aβ plaques are usually first deposited in the neocortex, before appearing in MTL regions, such as the EC and the HPC [4]. In our experiments we did not observe intraneuronal Aβ in the DG of 3xTg AD mice, and this contrasts with reports from the traditional perforant path projections between LEC layer II and the molecular/granular cell layers of the DG [56]. However, the DG is usually spared for neuropathological deposits in experimental models and patients until very late stages of AD [4, 24, 35, 57]. This provides an explanation as to why manipulation of activity levels in LEC layer II did not affect intraneuronal Aβ in the DG of 3xTg AD mice. Our findings of intraneuronal Aβ in CA1 can be explained by direct projections between LEC layer II and CA1, which bypasses the DG [58], the vulnerability of the EC-CA1 circuitry to neurodegeneration during AD [59, 60], or the first presentation of Aβ plaques appearing in CA1 of unmanipulated 3xTg AD mice [24].

Since aging is the major risk factor for developing sporadic AD, the disease pathology may be linked to the degradation of cellular clearance systems [11, 61, 62]. In line with this, previous reports suggest that deficits in the autophagic pathway are associated with AD progression and lead to an increase in intraneuronal Aβ aggregates [63, 64]. We observed lowered intraneuronal Aβ levels after LEC layer II neuronal silencing even in very young 3xTg AD mice, and this could also be explained by hyperactivity in this brain region early in the disease progression [65]. Neuronal hyperactivity in the HPC has been shown to induce Aβ pathology in mouse models [66, 67] and patients [68] with AD. In our experiments, the Sub was the first hippocampal subregion to display changes in intraneuronal Aβ after chronic LEC layer II silencing. The Sub is of high interest in the field, as it acts as a gateway between the neocortex and HPC [69, 70] and is affected early by Aβ pathology [71, 72]. Moreover, previous work has shown that lesions to the Sub resulted in reduced spreading of Aβ to interconnected regions in mice. However, the phasing of hippocampal subfield involvement in toxic intraneuronal Aβ build-up needs to be investigated further. In summary, we show that early intraneuronal Aβ levels in the LEC and HPC correlated with activity levels, and our findings give evidence to early AD neuropathology originating in select neuronal populations.

## 6. Acknowledgments

The authors would like to thank Dr. Rajeevkumar R. Nair for generating the viral vectors used in these experiments.

