## Supplementary material for "Manipulation of neuronal activity in the entorhinal-hippocampal circuit affects intraneuronal amyloid-β levels"

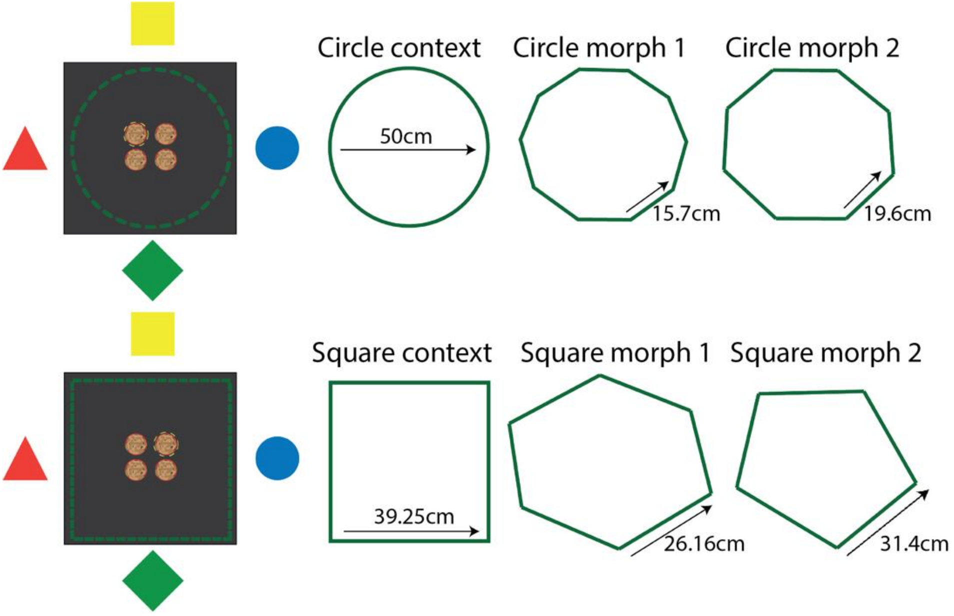


**Supplementary Figure 1. Context-dependent spatial memory task design.** Mice were initially taught to associate a specific reward location in a square- and a circle-chamber. The reward was buried in one of four cups with ginger-scented bedding. A trial was assessed as correct if the mice dug in the reward location associated with each chamber. If they passed the training phase (66.6% correct digging), their contextual memory performance was tested in the following morph chambers: a decagon (circle morph 1), an octagon (circle morph 2), a hexagon (square morph 1), and a pentagon (square morph 2). Mice were tested in the morph chambers for 4 days, with 8 sessions a day.

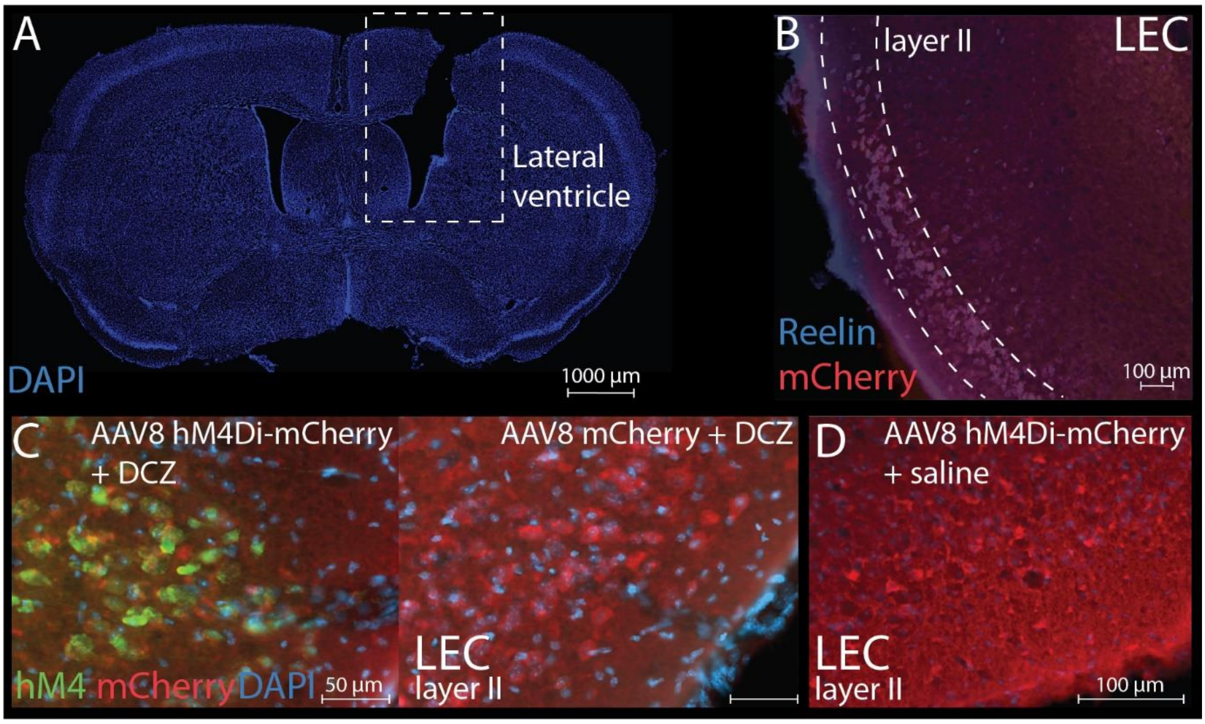


**Supplementary Figure 2. Verification of intraventricular cannula implantation and AAV-hM4D$_i$ injections in LEC layer II. A)** Intracranial cannula implantation site in the lateral ventricle of a 3xTg AD mouse visualized by DAPI nucleic staining (blue). The intracranial cannula was connected to an osmotic minipump via a catheter that allowed for the continuous and controlled infusion of either DCZ or saline for 2-3 weeks. **B)** Co-labelling of reelin (marker for principal fan cells) and mCherry following AAV-hM4D$_i$ injections into LEC layer II. **C)** *Left*: Expression of hM4 receptors (green) and mCherry (red) after AAV-hM4D$_i$ injections in LEC layer II and DCZ infusions. *Right*: Lack of hM4 receptor expression (green), but not mCherry (red) after AAV-mCherry injections in LEC layer II and DCZ infusions. **D)** Lack of hM4 receptor expression (green) after AAV-hM4D$_i$ injections (mCherry; red) in LEC layer II and saline infusions. Abbreviations; LEC: lateral entorhinal cortex; hM4D$_i$: modified inhibitory human muscarinic G-protein coupled receptor 4; hM4: human M4 muscarinic (hM4) inhibitory DREADDs; DCZ: deschloroclozapine.

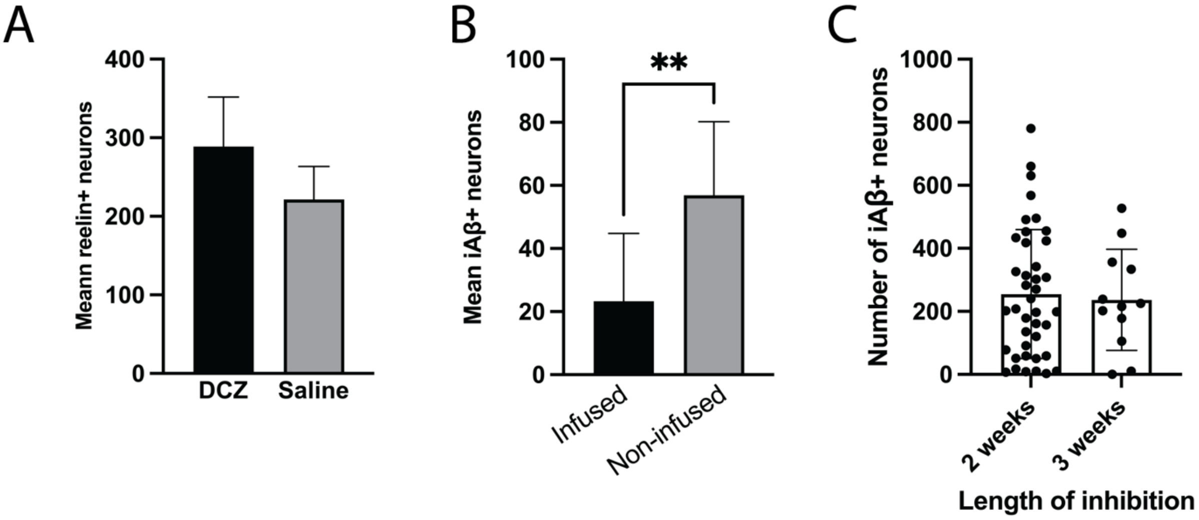


**Supplementary Figure 3. Supplementary effects of LEC layer II neuronal silencing in 3xTg AD mice. A)** Bar graph displaying mean intraneuronal reelin levels in LEC layer II neurons (<4 brain sections) after AAV-hM4D$_i$ injections and DCZ (*n* = 2) and saline (*n* = 2) infusions. **B)** Bar graph displaying mean intraneuronal Aβ+ LEC layer II neurons in infused and non-infused hemispheres (<4 brain sections) after AAV-hM4D$_i$ injections and DCZ (*n* = 25) infusions (*t* = 5.24, p < .01, unpaired two-tailed t-test). **C)** Box plots showing mean intraneuronal Aβ+ LEC layer II neurons after AAV-hM4D$_i$ injections and DCZ infusions for 2 (*n* = 21) or 3 (*n* = 4) weeks. Each dot represents a brain section containing the hippocampal subregions, and sections spanned the same rostro-caudal level for both AAV-hM4D$_i$ and AAV-mCherry injections. Abbreviations; iAβ: intraneuronal amyloid-β; DCZ: deschloroclozapine. Error bars denote ±1 SEM in panel A, and ±1 SD in panels B-C; **: p <.01.

### Supplementary Table 1: Antibodies used in experiments

| Antibody | Target | Identifier |
|---|---|---|
| Mouse anti-Aβ (McSA1) | Targets the N-terminal amino acids 1–12 of human Aβ | MediMabs Cat# MM-0015-1P, RRID:AB_1807985 |
| Mouse anti-reelin | Targets the extracellular matrix glycoprotein reelin | Millipore Cat# MAB5364, RRID:AB_2179313 |
| Mouse anti-mCherry | Targets mCherry fluorescence | Clontech Cat# 632543 |
| Rabbit anti-hM4 | Targets human muscarinic receptor 4 | Abcam Cat# ab189432 |
| Rabbit anti-cFos | Targets the cFos immediate-early gene | Abcam Cat# ab208942, RRID:AB_2747772 |
